# BDNF signaling requires Matrix Metalloproteinase-9 during structural synaptic plasticity

**DOI:** 10.1101/2023.12.08.569797

**Authors:** Diana Legutko, Bożena Kuźniewska, Katarzyna Kalita, Ryohei Yasuda, Leszek Kaczmarek, Piotr Michaluk

## Abstract

Synaptic plasticity underlies learning and memory processes as well as contributes, in its aberrant form, to neuropsychiatric disorders. One of its major forms is structural long-term potentiation (sLTP), an activity-dependent growth of dendritic spines that harbor excitatory synapses. The process depends on the release of brain-derived neurotrophic factor (BDNF), and activation of its receptor, TrkB. Matrix metalloproteinase-9 (MMP-9), an extracellular protease is essential for many forms of neuronal plasticity engaged in physiological as well as pathological processes. Here, we utilized two-photon microscopy and two-photon glutamate uncaging to demonstrate that MMP-9 activity is essential for sLTP and is rapidly (∼seconds) released from dendritic spines in response to synaptic stimulation. Moreover, we show that either chemical or genetic inhibition of MMP-9 impairs TrkB activation, as measured by fluorescence lifetime imaging microscopy of FRET sensor. Furthermore, we provide evidence for a cell-free cleavage of proBDNF into mature BDNF by MMP-9. Our findings point to the autocrine mechanism of action of MMP-9 through BDNF maturation and TrkB activation during sLTP.

## Introduction

Synaptic plasticity, the ability of synapses to modify their strength in response to their activity, is a fundamental process that underlies learning and memory. Long-term potentiation (LTP) is widely considered to be a cellular model of synaptic plasticity, and it has been extensively studied, predominantly on excitatory synapses located on small neuronal protrusions – dendritic spines (Nicoll, 2017). Synaptic strength corresponds, among other factors, to the number of AMPA (α-amino-3-hydroxy-5-methyl-4-isoxazolepropionic acid) receptors. After the LTP induction, AMPA receptors are rapidly inserted into the spine (Patterson *et al*, 2010; Penn *et al*, 2017; Shi *et al*, 1999) thus increasing the strength of a synaptic connection between two neurons. One of the manifestations of LTP is a structural change of dendritic spine morphology, where upon stimulation the spine increases its volume (Matsuzaki *et al*, 2004). To date, multiple molecular pathways have been identified that regulate synaptic plasticity (Hayashi, 2022), including a number of extracellular and transmembrane factors such as extracellular matrix (ECM), cell adhesion molecules, neurotrophic factors, and proteolytic enzymes that can process those molecules, with the matrix metalloproteinases (MMPs) serving as particularly prominent proteases (Wiera & Mozrzymas, 2021).

Among the MMPs, zinc-dependent endopeptidases, MMP-9 has emerged as a key player in regulating synaptic plasticity. Research on its involvement in LTP has been ongoing for almost two decades (Nagy *et al*, 2006; Okulski *et al*, 2007; Gorkiewicz *et al*, 2010, 2015; Lebida & Mozrzymas, 2017; Huntley, 2012; Vafadari et al., 2016). MMP-9 is considered one of the essential factors for the maintenance of LTP (Nagy *et al*, 2006) and is crucial for structural plasticity by contributing to a sustained increase of dendritic spines volume evoked by LTP (Wang *et al*, 2008). Upon stimulation, MMP-9 is released from a postsynaptic neuron to the extracellular space, where it can cleave several protein targets (Michaluk *et al*, 2007; Bajor *et al*, 2012; Wiera & Mozrzymas, 2021; Van Der Kooij *et al*, 2014; Ethell & Ethell, 2007; Peixoto *et al*, 2012). Notably, the proteolytic activity is tightly controlled by MMP-9 endogenous inhibitor, Tissue Inhibitor of Metalloproteinase-1 (TIMP-1) (Magnowska *et al*, 2016).

Several studies demonstrated that MMPs, including MMP-9, play a critical role in the maturation of brain-derived neurotrophic factor (BDNF) by processing its precursor form (proBDNF) to mature form (mBDNF) (Lee *et al*, 2001; Hwang *et al*, 2005; Mizoguchi *et al*, 2011; Niculescu *et al*, 2018). ProBDNF is a potent inhibitor of synaptic plasticity, while mBDNF is essential for maintenance of LTP (Kang *et al*, 1997; Lohof *et al*, 1993; Hedrick *et al*, 2016; Monteggia *et al*, 2004; Harward *et al*, 2016). mBDNF promotes synaptic plasticity through the activation of its receptor, tropomyosin receptor kinase B (TrkB) that triggers multiple downstream signaling pathways, including the mitogen-activated protein kinase (MAPK), phospholipase Cγ (PLCγ) and phosphoinositide 3-kinase (PI3K) pathways that are known to be pivotal for the induction and maintenance of synaptic plasticity (Cunha *et al*, 2010).

Despite significant progress in our understanding of the role of MMP-9 in synaptic plasticity, a number of questions remain unanswered. In particular, the kinetics of MMP-9 release and proteolytic action, especially in conjunction with proBDNF processing, is unknown. Depending on the report and protocol used, MMP-9 release has been shown to be as early as 5 minutes upon neuronal stimulation (Michaluk *et al*, 2007), through 15–30 minutes (Nagy *et al*, 2006), and up to 1 hour (Nagy *et al*, 2006, 2007; Bozdagi *et al*, 2007). In the studies where MMP-9 was applied externally as a recombinant protein, it was usually incubated for either 30 (Michaluk *et al*, 2011; Wang *et al*, 2008) or 40 minutes with the cultured neurons (Magnowska *et al*, 2016) to cause morphological change of dendritic spines. In comparison, BDNF, which can be released in pro- and mature-form, has been shown to be secreted rapidly (within seconds) upon increase in K^+^ concentration (Dean *et al*, 2009), theta-burst stimulation (Matsuda *et al*, 2009) or sLTP-inducing glutamate uncaging (Harward *et al*, 2016). Additionally, it has been shown that TrkB activation is also almost immediate, reaching the maximum within 1 minute from the onset of stimulation (Harward *et al*, 2016). If MMP-9 activity indeed causes maturation of proBDNF and leads to TrkB activation (Hwang *et al*, 2005; Mizoguchi *et al*, 2011; Niculescu *et al*, 2018), both proteins must coincide in space and time around a dendritic spine. This hypothesis could address a long-standing debate about which enzymes process proBDNF to mBDNF and in which compartments it occurs (Sasi *et al*, 2017). Hence, it is essential to understand the spatiotemporal dynamics of MMP-9 action at the level of single dendritic spines.

This report provides insights into the kinetics of MMP-9 release and activation of TrkB receptors during LTP-evoked structural plasticity. In order to pinpoint the dynamic processes of structural plasticity, protein release, and receptor activation, we have used two-photon imaging, 2-photon fluorescence lifetime imaging microscopy (2pFLIM) to measure Förster Resonance Energy Transfer (FRET) sensor of TrkB, and 2-photon glutamate uncaging on individual dendritic spines (Colgan *et al*, 2018; Tu *et al*, 2020; Chang *et al*, 2019; Harward *et al*, 2016). These techniques allowed us to test how MMP-9 influences structural LTP (sLTP) evoked by glutamate uncaging. We have observed MMP-9 release upon sLTP and its effect on TrkB activation. Moreover, in a cell-free assay, we have confirmed proBDNF cleavage to mBDNF by MMP-9. Overall, our results indicate that both MMP-9 and BDNF are released from stimulated spines, leading to autocrine TrkB activation, and support the hypothesis that rapid maturation of BDNF may occur in the extracellular space upon stimulation.

## Results

### Spine-head enlargement during sLTP depends on MMP-9 activity

Previous studies have shown that dendritic spine volume and shape correlates with the size of the synapse (Borczyk *et al*, 2019; Vardalaki *et al*, 2022), number of synaptic AMPA-type glutamate receptors (Matsuzaki *et al*, 2001; Shi *et al*, 1999). Therefore, in this study, we focused on dendritic spine volume as a proxy of synaptic plasticity (Tønnesen & Nägerl, 2016). Pyramidal neurons of CA1 subfield of the hippocampus were transfected in organotypic culture using a biolistic method with a plasmid encoding GFP. Successfully transfected neurons were imaged using two-photon microscope. During sLTP stimulation (see Methods and Harward *et al*, 2016), uncaging of 4-methoxy-7-nitroindolinyl (MNI)–glutamate caused rapid and transient three-to four-fold spine volume increase, which subsequently stabilizes at an elevated level, typically ranging from 150-200% of its original size. This sustained phase starts around 8 minutes after stimulation (Fig. 1A-C, Supplementary Video 1). Similarly to previous studies, the spine volume change was assessed as a ΔF/F_0_ of GFP fluorescence for individual spines in time (Harward *et al*, 2016; Hedrick *et al*, 2016) (see Methods).

**Figure 1.**
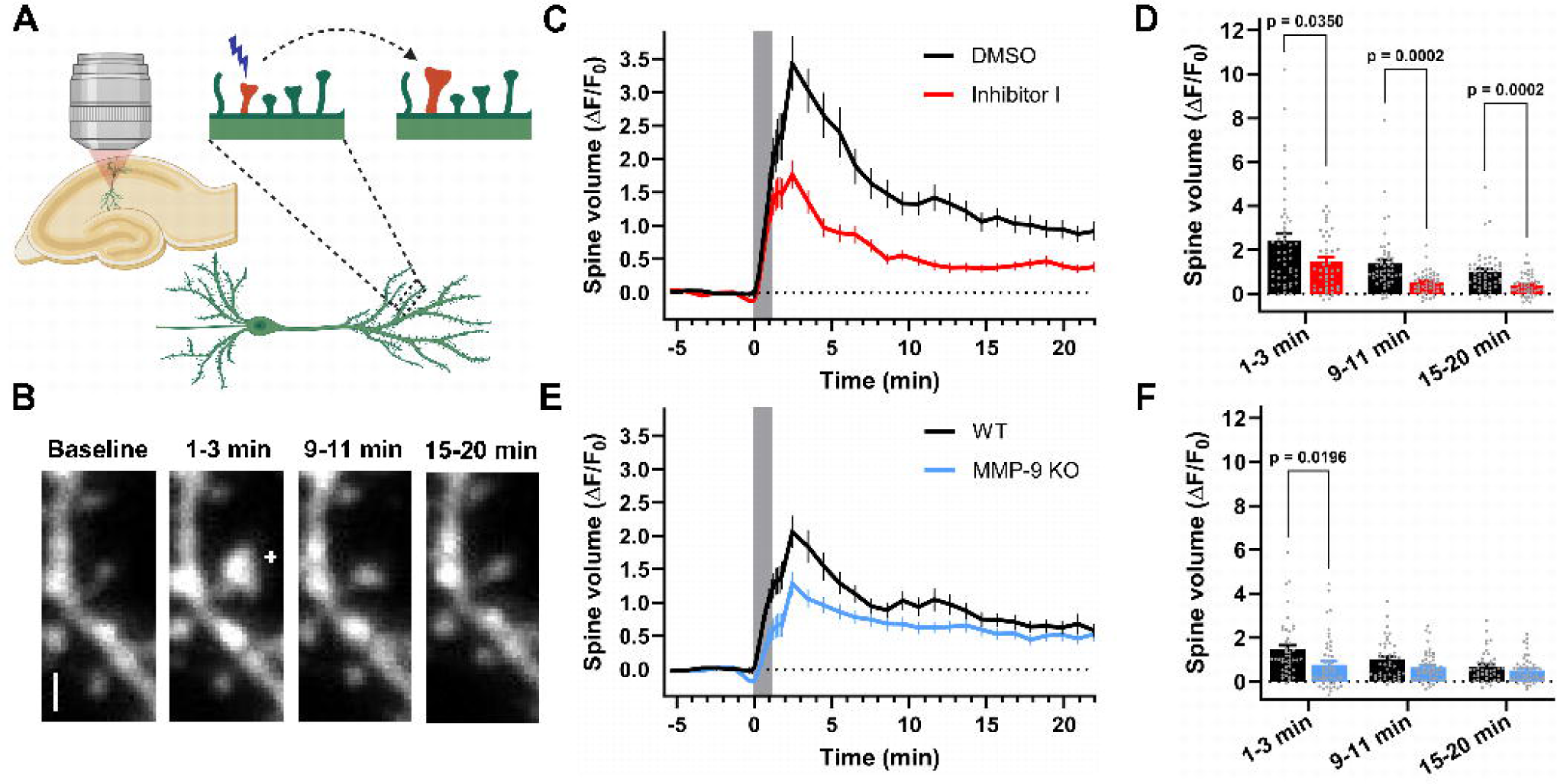
Spine-head enlargement during sLTP depends on MMP-9 activity. **A** Experimental arrangement: GFP-expressing neuron in CA1 subfield of hippocampus. Individual spine in secondary dendrite was stimulated using sLTP protocol leading to the spine enlargement. **B** Time-lapse z-integrated (z-stack) images of spines stimulated at time 0 min by sLTP protocol. Cross indicate a spot of two-photon uncaging, scale bar: 1 μm. **C** Averaged time courses of changes in spine volume (measured as ΔF/F_0_) for spines stimulated by uncaging. Data are means ± SEM, DMSO (n = 50 spines; 27 cells, 15 animals), Inhibitor I (n = 46 spines; 21 cells, 10 animals). Grey box indicates duration of sLTP protocol. **D** Statistical analysis of (C); Averaged spine volume changes for three phases: 1-3 min. (transient), 9-11 and 15-20 min. (sustained). Gray dots represent individual values for spines, bars are means ± SEM (black – DMSO, red – Inhibitor I). Repeated measures ANOVA (Time (F (1.175, 110.5) = 33.60, p < 0.0001); Inhibitor (F (1, 94) = 18.63, p < 0.0001); TimeˣInhibitor (F (2, 188) = 0.6494, p = 0.5235)) followed by Šidák’s multiple comparison test (DMSO vs Inhibitor I at 1-3 min (p = 0.0350, 95% C.I. = [0.05124 to 1.863], 9-11 min (p = 0.0002, 95% C.I. = [0.3747 to 1.358]) and at 15-20 min (p = 0.0002, 95% C.I. = [0.2553 to 0.9569])). **E** as in (C), but for WT (n = 44 spines; 13 cells, 7 animals) and MMP-9 KO animals (n = 51 spines; 14 cells, 7 animals). **F** Statistical analysis of (E), marks as in (D). Bars are means ± SEM (black – WT, blue – MMP-9 KO). Repeated measures ANOVA (Time (F (1.463, 136.1) = 12.56, p < 0.0001); MMP-9 KO (F (1, 93) = 8.444, p = 0.0046); TimeˣMMP-9 KO (F (2, 186) = 3.253, p = 0.0409) followed by Šidák’s multiple comparison test (WT vs MMP-9 KO at 1-3 min (p = 0.0196, 95% C.I. = [0.08769 to 1.311]).

To test if MMP-9 activity is necessary for sLTP, we first used a chemical inhibitor of MMP-9 and MMP-13 (Inhibitor I, 5 μM; Fig. 1C) and compared the effects with the vehicle (DMSO) control. We assessed spine head volume at three time intervals of sLTP: the transient phase (1-3 minutes after the onset of uncaging), and the sustained phase (9-11 minutes and 15-20 minutes after the uncaging). Presence of Inhibitor I significantly diminished spine volume increase in all phases (Fig. 1C, 1D). Therefore, MMPs (including MMP-9) activity appears to be important both during the induction and maintenance of sLTP (as described before).

There are over twenty MMPs with overlapping substrate specificity (Fields, 2015; Cieplak & Strongin, 2017) and there are no fully specific, commercially available inhibitors for MMP-9. Therefore, to further test the involvement of the protease in sLTP, we have used hippocampal slice cultures prepared from MMP-9 KO mice and their WT littermates (Fig. 1E, 1F). Unlike chemical inhibitors, MMP-9 KO significantly reduced spine volume change only during the transient phase of sLTP (1-3 min, Fig. 1F), but not during the sustained phase. These results suggest that MMP-9 acts rapidly (within 1-3 min) on biochemical processes leading to sLTP.

### MMP-9 release during sLTP

The proteolytic activity of MMP-9 in neuronal tissue is tightly controlled on many levels, from transcription, through local translation, release, and inhibition by TIMP-1 (Michaluk *et al*, 2007; Dziembowska *et al*, 2012; Magnowska *et al*, 2016; Szklarczyk *et al*, 2002; Vafadari et al., 2016). However, the exact moment of MMP-9 secretion is unknown during LTP. Our previously published results have shown an increase of MMP-9 level in extracellular neuronal culture medium 5 minutes post neuronal stimulation (with no earlier time-points investigated, see: Michaluk *et al*, 2007; Stawarski *et al*, 2014), yet many studies indicate that mBDNF has to be present during LTP induction to maintain plasticity (Pang *et al*, 2016; Harward *et al*, 2016). If the molecular mechanism of MMP-9 signaling affects the maturation of proBDNF and activation of TrkB, MMP-9 secretion has to be as fast as the release kinetics reported for BDNF (Dean *et al*, 2009; Matsuda *et al*, 2009; Harward *et al*, 2016). To verify if MMP-9 is released from dendritic spines in an activity-dependent manner, we developed pH-sensitive green fluorescent protein (superecliptic pHluorin, SEP) (Miesenböck *et al*, 1998) fused to C-terminus of hemopexin domain of proteolytically inactive mouse MMP-9. Cells in hippocampal organotypic cultures were co-transfected with the MMP9-SEP and mCherry (the latter for the visualization of dendritic spine morphology; Fig. 2A-B, Supplementary Video 2). To study MMP-9 release during sLTP, we imaged dendritic spines of CA1 pyramidal neurons with 2-photon microscopy at 7.8 frames per second, while stimulating neurons with glutamate uncaging (0.49 Hz or 16 frames, 30 times).

**Figure 2.**
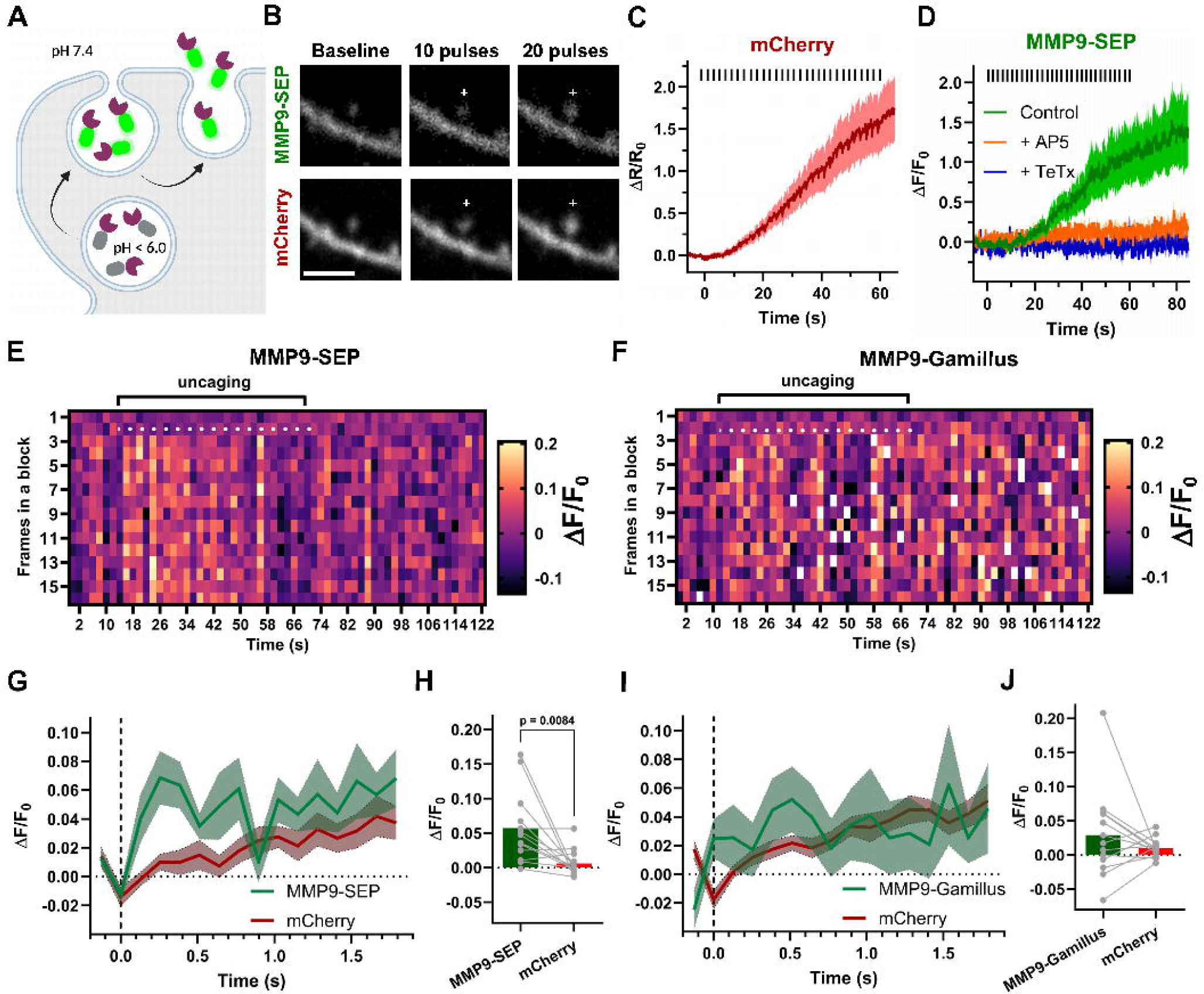
MMP-9 release during sLTP. **A** MMP9-SEP fluorescence is quenched by low (< 6.0) pH inside vesicles. Stimulation of the spine promotes exocytosis. Once the vesicle fuses with the cell membrane, pore opens and the acidic environment in the vesicle’s lumen is neutralized (pH 7.4) allowing for observation of fluorescence of SEP. **B** Time-lapse t-integrated (t-stack) images of spines stimulated by sLTP protocol – average of 10 consecutive frames collected after indicated uncaging pulses. Separate channels of MMP9-SEP and mCherry are shown. Crosses indicate a spot of two-photon uncaging, scale bar: 1 μm. **C** Averaged time courses of changes in volume (measured as ΔR/R_0_ for mCherry channel) for spines stimulated by uncaging. Data are means ± SEM (n = 13 spines, 3 cells, 1 animal), timing of glutamate pulses during sLTP protocol are indicated by black bars. **D** Averaged time courses of changes in SEP fluorescence (measured as ΔF/F_0_) for spines stimulated by uncaging. Data are means ± SEM (Control n = 13 spines, 3 cells, 1 animal; + AP5 n = 13 spines, 3 cells, 2 animals; + TeTx n = 12 spines, 1 cell, 1 animal), timing of glutamate uncaging laser pulses during sLTP protocol are indicated by black bars. **E** Heatmap representing averaged fluorescence of MMP9-SEP in stimulated spines (n=13). Each colored rectangle represents relative fluorescence intensity of SEP. Imaging data were divided into blocks (columns) of 16 frames matching uncaging frequency, so that uncaging pulse always occurs in a second frame of the block. sLTP protocol starts after recording baseline of 7 blocks (∼ 14 s) and lasts for 60 seconds (30 pulses, 0.5 Hz). White dots represent uncaging laser pulse which occurs always during the second frame in a block during sLTP-evoking protocol. **F** same as (E) but for MMP9-Gamillus. **G** Averaged time courses of changes in MMP9-SEP and corresponding mCherry fluorescence (measured as ΔF/F_0_) for spines stimulated by uncaging. The plot represents superimposed and averaged blocks of 16 frames (2.048 s) for normalized fluorescence from 2^nd^ to 18^th^ uncaging pulse. Data are means ± SEM (n = 13 spines, 3 cells, 1 animal), timing of glutamate uncaging laser pulses during sLTP protocol are indicated by dashed vertical line at time = 0 s. **H** Peak of the uncaging-triggered change in MMP9-SEP and mCherry fluorescence (calculated as a mean of ΔF/F_0_ for the three subsequent frames after uncaging pulse: 0.128-0.384 s). Dots with lines represent values for individual spines for MMP9-SEP and mCherry fluorescence paired together. Bars are means. Paired t-test (t=3.149, df=12, p = 0.0084;). **I** Same as in (G) but for MMP9-Gamillus (n = 13 spines, 5 cells, 2 animals). **J** same as in (H) but for MMP9-Gamillus. Paired t-test (t=1.024, df=12, p = 0.3262).

MNI-glutamate uncaging caused an increase in mCherry fluorescence, indicative of spine volume enlargement during sLTP protocol (Fig. 2C). Observation of corresponding MMP-9-SEP fluorescence showed an increase in signal restricted to the stimulated spine (Fig. 2B, 2D), suggesting the release of MMP-9. In order to explore if the MMP-9 release is NMDAR (N-methyl-D-aspartate receptor)-dependent, we applied sLTP protocol in the presence of NMDAR antagonist AP5. This manipulation abolished SEP fluorescence increases during stimulation. Similarly, when the cells were co-transfected with tetanus toxin light chain (TeTx), which blocks the synaptobrevin-dependent exocytosis (Link *et al*, 1992; Harward *et al*, 2016), we did not observe the SEP fluorescence increases during sLTP protocol (Fig. 2D). Therefore, we concluded that synaptic release of MMP-9 requires its exocytosis and is NMDAR-dependent.

To further assess the fast kinetics of MMP9 releases after each uncaging pulse, we divided imaging data into blocks of 16 frames (corresponding to uncaging intervals, 2.048 s), and stacked them in such a way that the uncaging pulse was always applied during the second row of the block (Fig. 2E). Then, the fluorescence data were normalized to the first two averaged rows in each block. This procedure allowed us to visualize the averaged fluorescence changes normalized to fluorescence before uncaging (averaged over 2 frames) as a heatmap (Fig. 2E). The resulting image suggests that fluorescence increases appear after the second uncaging pulse and diminishes after the 22^nd^ uncaging pulse (Fig. 2E).

In order to study if the increases in MMP9-SEP fluorescence were a result of pH changes associated with MMP9 release, we performed the same experiments using an analogous fusion protein to MMP9-SEP, but with pH-stable form of GFP, Gamillus (Shinoda *et al*, 2018). We did not observe a fluorescence increase in the heatmap from MMP9-Gamillus (Fig. 2F).

To better visualize the rapid release of MMP-9 upon uncaging, we have averaged relative fluorescence for the 2-18 uncaging pulses and plotted them as a function of time, together with the corresponding fluorescence of mCherry (Fig. 2G, 2H). In the mCherry channel, we observed a slow fluorescence increase corresponding to spine volume change. MMP9-SEP fluorescence showed a much faster onset, suggesting a rapid release of MMP9 in response to each uncaging pulse. MMP9-Gamillus, however, showed signals similar to mCherry (Fig. 2I, 2J). Therefore, these results further support a fast release of MMP-9 at the synapses undergoing sLTP.

### TrkB activation depends on MMP-9 activity

Previous studies suggested proBDNF to be a potential target for MMP-9 proteolytic activity that leads to maturation of the neurotrophic factor and activation of TrkB (Hwang *et al*, 2005; Martinelli *et al*, 2021; Niculescu *et al*, 2018; Kaminari *et al*, 2017; Mizoguchi *et al*, 2011). Given that our and other results indicate that both BDNF and MMP-9 are rapidly released upon synaptic stimulation, we hypothesized that MMP-9 took part in the maturation of BDNF. To test this hypothesis, we transfected neurons in hippocampal organotypic cultures with a TrkB FRET sensor made of TrkB fused with GFP and SH2-domain of PLC-delta fused with a pair of mCherry proteins (Harward *et al*, 2016). Then, we measured TrkB activation (Harward *et al*, 2016) during uncaging-evoked sLTP (Fig. 3B, Supplementary Video 3) in the presence of either Inhibitor I or DMSO for control (Fig. 3C, 3D). Inducing sLTP with glutamate uncaging decreased fluorescence lifetime of GFP and increased the calculated binding fraction between TrkB and SH2-domain of the sensor (Fig. 3A, 3C), indicating the activation of TrkB (Fig. 3B). Inhibitor I significantly attenuated TrkB activation in comparison to DMSO after the sLTP induction in both transient phase and sustained phases of sLTP (Fig. 3D) indicating that MMP-9 activity is most probably involved in TrkB activation.

**Figure 3.**
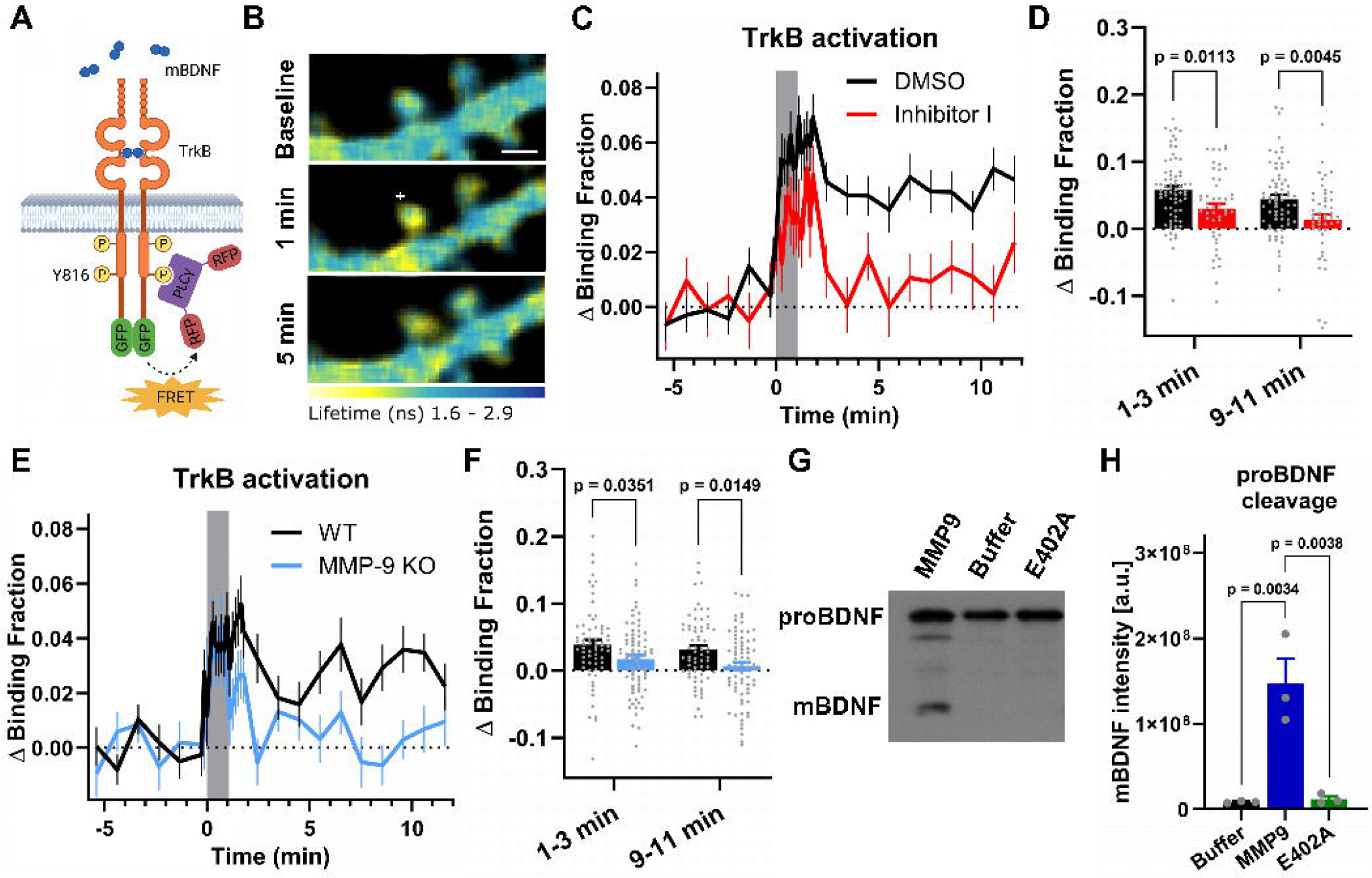
TrkB activation depends on MMP-9 activity. **A** TrkB FRET sensor schematic. **B** Two-photon FLIM images of TrkB activation averaged at indicated time points. Warmer colors represent shorter lifetimes, increased binding fraction and higher TrkB activity. White cross indicate uncaging spot, scale bar: 1 μm. **C** Averaged time courses of TrkB activation (measured as a change of the sensor binding fraction) for spines stimulated by uncaging. Data are means ± SEM. Grey box indicates duration of sLTP protocol. **D** Statistical analysis of (C). Averaged TrkB activation (measured as a Δ Binding Fraction) for transient phase (1-3 min after sLTP induction) and sustained phase (9-11 min. after sLTP induction) in spines incubated with DMSO (n = 70 spines, 29 cells, 16 animals) or Inhibitor I (n = 49 spines, 21 cells, 10 animals). Gray dots represent individual values for spines, bars (black – DMSO, red – Inhibitor I) are means ± SEM. Repeated measures ANOVA (Time (F (1, 117) = 11.46, p = 0.0010); Inhibitor (F (1, 117) = 10.80, p = 0.0013); TimeˣInhibitor (F (1, 117) = 0.1103, p = 0.7403)) followed by Šidák’s multiple comparison test (DMSO vs Inhibitor I at 1-3 min (p = 0.0113, 95% C.I. = [0.005425 to 0.05049]) and at 9-11 min (p = 0.0045, 95% C.I. = [0.008394 to 0.05346])). **E** Same as in (C). Averaged time courses of TrkB activation observed in spines obtained from WT, and MMP-9 KO mice. **F** same as in (D). WT (black bars, n = 66 spines; 22 cells, 10 animals), MMP-9 KO (blue bars, n = 73 spines; 25 cells, 11 animals). Repeated measures ANOVA (Time F (1, 137) = 5.084, p = 0.0257; MMP-9 KO F (1, 137) = 7.982, p = 0.0054; TimeˣMMP-9 KO F (1, 137) = 0.1253, p = 0.7238)) followed by Šidák’s multiple comparison test (WT vs MMP-9 KO at 1-3 min (p = 0.0351, 95% C.I.=[0.001241 to 0.04204] and at 9-11 min (p = 0.0149, 95% C.I. = [0.004051 to 0.04485])). **G** Representative immunoblot of digestion reaction of proBDNF incubated with active MMP-9, inactive MMP-9 (E402A), or only in the reaction buffer. Arrows indicate proBDNF (∼26 kDa) and mBDNF (∼14 kDa). **H** Quantification of three separate Western-blots of proBDNF digestion. Gray dots represent individual values of mBDNF band intensity in separate experiments and Western-blots for each experimental condition. Bars are means ± SEM. One-way ANOVA (F (2, 6) = 20.38, p = 0.0021) followed by Tukey’s multiple comparisons test (Buffer vs MMP-9 p = 0.0034, 95% C.I. = [-213821761 to -62305372] and E402A vs MMP-9 p = 0.0038, 95% C.I. = [59136932 to 210653321]).

To further test if MMP-9 is necessary for TrkB activation, we used hippocampal organotypic cultures prepared from MMP-9 KO mice and their WT littermates (Fig. 3E), and measured TrkB activation with the TrkB sensor. Similarly to the application of Inhibitor I, MMP-9 KO influenced TrkB activity during both transient and sustained phases of structural plasticity. These results demonstrate that MMP-9 is required for both transient and sustained activation of BDNF receptor following repeated glutamate uncaging evoking sLTP.

Finally, we examined if MMP-9 can cleave proBDNF into mBDNF in a cell-free assay. To do this, we incubated a recombinant proBDNF with recombinant active MMP-9 overnight at 37°C in a reaction buffer (see Methods). As a control, we used recombinant inactive mutant (E402A) of MMP-9 or a buffer without MMP-9 (Michaluk *et al*, 2011). The cleavage of proBDNF was quantified by Western blot, using anti-BDNF antibody recognizing both (pro- and mature-form). Incubation with active MMP-9, but not with mutant MMP-9, causes cleavage of proBDNF (Fig. 3G). These results indicate that MMP-9 cleaves proBDNF into mBDNF.

## Discussion

In this report, we demonstrate that: (i) MMP-9 inhibition impairs structural synaptic plasticity (sLTP) evoked by repetitive stimulation by glutamate uncaging, (ii) MMP-9 is rapidly, within seconds, released from the dendritic spines, (iii) MMP-9 inhibition impairs activation of BDNF receptor TrkB, and (iv) MMP-9 partially cleaves proBDNF to produce its mature form, known to activate TrkB and evoke structural synaptic plasticity. In aggregate, these results demonstrate a mechanistic link between MMP-9 and BDNF in controlling synaptic plasticity.

The MMP-9 effects were evident during sustained phase of sLTP (9-11 or 15-20 min post-stimulation), and even more robustly during the transient phase (1-3 min post-stimulation). The application of MMP-9/-13 Inhibitor I and its effect on sLTP at the sustained phase of sLTP is in agreement with previously published electrophysiology and imaging data (Nagy *et al*, 2006; Wang *et al*, 2008; Gorkiewicz *et al*, 2015; Wiera *et al*, 2013). However, in slice cultures prepared from MMP-9 KO animals, we did not observe the influence of knockout on the sustained phase of sLTP. It is possible that MMP-9 KO mice have some compensatory mechanisms and lack of MMP-9 is compensated for example by activity of another MMP. Additionally, even though MMP-9 KO mice are kept on C57BL/6 background and the line is maintained by breeding heterozygotes together, WT littermates which have been used in the study have markedly lower increase of spine volume in comparison to slices prepared from wildtype C57BL/6J mice (Fig. 1). Therefore, the difference in sLTP phenotypes may be explained by some unaccounted genetic differences between the two mouse lines. It has been recently shown that even closely-related mice substrains C57BL/6J and C57BL/6N express substantial metabolic and genetic differences (Nemoto *et al*, 2022).

This early signaling by MMP-9 during sLTP apparently contradicts with previous studies suggesting that MMP-9 is required for late-phase LTP, but not for early-phase LTP (Nagy *et al*, 2006; Gorkiewicz *et al*, 2015; Wiera *et al*, 2017). Interestingly, in the paper of Nagy *et al*, (2006), LTP in MMP-9 KO animals are impaired in both early- and late-LTP, but the use of chemical inhibitors of MMP-9 affected only late-LTP. While both Gorkiewicz *et al*. (2015) and Wiera *et al*. (2013, 2017) reported impairment of only late-LTP, the study of Gorkiewicz et al. was done in amygdala and Wiera *et al*.(2013) on mossy fiber-CA3 pathway where LTP is mostly presynaptic and independent on NMDA receptors (Nicoll & Schmitz, 2005). Their observed effects of MMP-9 KO on sLTP are in agreement with our data showing a rapid MMP-9 release upon synaptic stimulation (Fig. 2) and fast activation of TrkB. Both these effects might be explained by the release of the enzyme which was already accumulated at the postsynaptic site. This hypothesis may appear contradictory to the previous findings that MMP-9 is locally translated in response to synaptic stimulation and then released to contribute to the synaptic plasticity, the processes that require minutes to occur (Dziembowska *et al*, 2012; Michaluk *et al*, 2007; Stawarski *et al*, 2014). It should be, however, noted that there have been previous reports showing an expression of MMP-9 protein within the dendritic spines under the basal condition (Wilczynski *et al*, 2008; Michaluk *et al*, 2007; Aujla & Huntley, 2014). Interestingly, MMP-9 protein was predominantly found in small dendritic spines which are more prone for plastic changes (Kasai *et al*, 2010). Additionally, differences in LTP-evoking protocols might also explain why the effect of MMP-9 inhibition on the transient phase of sLTP has not been observed before. Wang and co-workers (2008) have shown that MMP-9 is both necessary and sufficient to drive spine enlargement and synaptic potentiation concomitantly by using theta-stimulation-induced LTP paired with a depolarization of postsynaptic neurons. This pairing protocol, due to backpropagating action potential and presynaptic activation, can itself lead to the postsynaptic or presynaptic exocytotic release of BDNF and MMP-9. Therefore the glutamate action on the postsynaptic receptors in releasing those molecules could not be defined. In contrast, our single spine sLTP protocol provided direct evidence that BDNF and MMP-9 plays a role in spine-specific signaling events. Moreover, a recent study of Stein *et al*. (2021) dissociated the NMDAR-dependent structural and functional LTP by pointing to the role of non-ionotropic NMDAR signaling through p38 MAPK. This report might further explain the differences in the kinetics of MMP-9 involvement in electrophysiological and uncaging LTP experiments.

Our results showed that MMP-9 can be rapidly released upon stimulation, which further supports the early action of this protease for sLTP induction. Importantly, we compared changes in fluorescence intensity of MMP9-SEP and mCherry (morphological marker) in activated spines between uncaging pulses, and these data indicate that spine volume increase alone cannot explain the observed changes in SEP fluorescence. Therefore, we infer that observed fluorescence increase is related to exocytosis events in the dendritic spine. Presented results suggest that MMP-9 release may occur within 2 seconds after the first uncaging pulse, remain sustained over 40 seconds, followed by a decrease to the baseline level, which may be due to the depletion of spine vesicles. The timescale of MMP-9 release is thus similar to that of BDNF release (Harward *et al*, 2016) and AMPA receptors (Patterson *et al*, 2010), suggesting that the initial phase of sLTP involves a rapid increase in various exocytosis activity.

To date, there have not been many identified ECM substrates of MMP-9 in the brain. On the other hand, post-synaptically-originating or trans-synaptic cell adhesion molecules, such as ICAM-5 (Conant *et al*, 2010), β-dystroglycan (Michaluk *et al*, 2007), nectin-3 (Van Der Kooij *et al*, 2014), CD44 (Bijata *et al*, 2017) or neuroligin-1 (Peixoto *et al*, 2012) have been shown to be partially cleaved by MMP-9. Similarly, MMP-9 has been shown to be involved in cleavage of proBDNF into its mature form (mBDNF) (Hwang *et al*, 2005; Mizoguchi *et al*, 2011). Mizoguchi et al. (2011) has shown that pentylenetetrazole kindling of mice over several days increases a proportion of mature to proBDNF, and this effect was diminished in MMP-9 KO animals. Additionally, Hwang et al. (2005) incubated membrane fraction of cortical culture overexpressing proBDNF with high concentration of recombinant MMP-9 and observed proBDNF cleavage to mBDNF. The aforementioned results could not ascertain direct pro-BDNF cleavage by MMP-9 as opposed to the participation of the enzyme in a proteolytic cascade. To complement the existing knowledge, we showed directly the cleavage of proBDNF to mBDNF by MMP-9 without other proteases that could participate in the process or without the presence of TIMPs in the samples which might interfere with cleavage by MMPs. The presented results further support the MMP-9 involvement on BDNF maturation by demonstrating a detailed time-course of MMP-9 activity at the excitatory synapse and the contribution of this protease to TrkB activation. Thus, our study gives insights into the ongoing discussion whether BDNF is released from dendrites in its mature- or pro-form (Sasi *et al*, 2017).

Our results of TrkB activation may suggest mixed nature of its ligand. Decreased MMP-9 activity either by application of chemical inhibitor or gene knockout leads to a decreased level of TrkB activation during the transient phase of sLTP. Furthermore, while under the control condition, the receptor activation is sustained (∼ 10 min. after activation), under an MMP-9 inhibition, it reverses back to the baseline. It is possible that vesicles containing mBDNF are being depleted first and the mature ligand is acting directly during stimulation phase on TrkB (Matsumoto *et al*, 2008). In contrast, vesicles containing unprocessed proBDNF could serve as reservoir of the ligand that requires time to be activated by MMP-9. This might also explain why previous reports have found MMP-9 role in late-rather than early-LTP.

Given that the release kinetics of BDNF and MMP-9 are similar, one could speculate that the effect of MMP-9 inhibition on early TrkB activation can be achieved because both, MMP-9 and BDNF are co-localized and co-released from the same release vesicles. This would provide spatiotemporal coordination for proBDNF cleavage by MMP-9.

Overall, our data provide insights into the pivotal role of MMP-9 in synaptic plasticity and associated TrkB activation, demonstrating rapid (seconds) MMP-9 availability and activity. In addition, we provide a molecular link between the activity of the protease and function of BDNF, by following the activity of its receptor, TrkB.

## Methods

### Animals

All animal procedures were approved by the Max Planck Florida Institute for Neuroscience, and Use Committees and were conducted in accordance with the NIH Guide for the Care and Use of Laboratory Animals, as well as with the Animal Protection Act of Poland (directive 2010/63/EU). In this study we used 4-8 days-old mice of either sex. C57BL/6J mice were obtained from Charles River and MMP-9 KO ( B6.FVB(Cg)-Mmp9tm1Tvu/J ) were purchased from Jackson Laboratory. MMP-9 KO mice has been maintain by mating heterozygotes with heterozygotes. The genotype of each animal used was verified before preparing slices using PCR of genomic DNA isolated from the tail. For experiments we used MMP-9 KO mice and their wild-type (WT) littermates.

### Plasmids

pCAG_GFP, pCAG_mCherry, pCAG_MMP-9 – full length mouse (MMP-9) from pcDNA3.1_MMP-9-HA (Addgene #121172) cloned into pCAG vector, pCAG_MMP-9a1205c -full length mouse MMP-9 with point mutation in single nucleotide 1205 from adenine to cytosine resulting in codon change E402A which results in lack of MMP-9 enzymatic activity. Primers used for mutagenesis: Fwd: 5’-TGGCAGCGCACGCGTTCGGCCATGC, Rev: 5’-GCATGGCCGAACGCGTGCGCTGCCA, primers used for NEBuilder HiFi DNA Assembly: Fwd: 5; tgtctcatcattttggcaagGTGGTGGAATTCATGAGTC, Rev: 5’ tgctcaccatAGCGTAATCTGGAACATC. pCAG_MMP-9a1205c-SEP, SEP obtained by PCR reaction and cloned into pCAG_MMP-9a1205c vector with a use of NEBuilder HiFi DNA Assembly. Pimers: Fwd: 5’ CCATACGATGTTCCAGATTACGCTATGAGTAAAGGAGAAGAACTTTTCACTGG, Rev: 5’ GTTTAAACGGGCCCTCTAGACTCGAGCGGCCGCTTATTTGTATAGTTCATC For pCAG_MMP9a1205c-Gamillus, Gamillus obtained by PCR reaction from (Addgene #124837) was inserted in place of SEP with a use of NEBuilder HiFi DNA Assembly. Primers: Fwd: 5’-agattacgctATGGTGAGCAAGGGCGAG, Rev: 5’: ggcagagggaaaaagatccgGCAGAATTCTTACTTGTACAGCTCG. pCAG_mCherry-Tetx obtained by Yasuda Laboratory^21^. pCAG_TrkB-GFP –Tropomyosin receptor kinase B conjugated with GFP at C terminus acting as FRET (Förster resonance energy transfer) donor for PLC RFP, pCAG_PLC-RFP – SH2 domain of phospholipase Cγ1 (PLC-γ1) conjugated with two mRFP1 at its N and C terminus acting as acceptor for TrkB FRET sensor.

### Recombinant, inactive MMP-9 (E402A) production

Expression of the previously described(Michaluk *et al*, 2011) recombinant non-active mutant of MMP-9(E402A) was performed using the Bac-to-Bac Baculovirus expression system, according to the manufacturer’s instructions (Thermo Fisher Scientific). Briefly, DH10Bac competent cells, were transformed with pFastBac1_MMP-9(E402A) mutant was transform. Colonies that performed transposition of recombinant plasmid fragment into bacmid DNA were identified by blue–white selection, and recombinant bacmid was isolated and verified by PCR. The Sf21 insect cells were transfected with recombinant bacmid using Cellfectin reagent (Thermo Fisher Scientific) to obtain recombinant baculovirus. After amplification and titration of the recombinant baculovirus, High-Five cells were infected and incubated in the Sf-900IISFM serum-free medium (Thermo Fisher Scientific). Conditioned medium was collected 48 hours after infected, medium was collected and MMP-9(E402A) was purified by affinity chromatography with gelatin–Sepharose 4B (Cytiva) as previously described(Sadatmansoori *et al*, 2001). Protein concentrations in the collected fractions were measured using Bradford reagent (Sigma).

### proBDNF digestion assay

Twenty ng of recombinant proBDNF (Alomone Labs) was incubated with 50 ng of recombinant MMP-9 (Calbiochem) or 50 ng of recombinant, human, inactive MMP-9(E402A) in total volume of 20 μl. All reactions were incubated in reaction buffer (final concentration: 150 mM NaCl, 10 mM CaCl_2_, 0.01% BSA, 50 mM Tris-Cl, pH7.5) for 16 hours at 30°C.

### Western blotting

ProBDNF digestion samples were run on 15% SDS-PAGE gels and electro-transferred onto polyvinylidene difluoride membrane (Immobilon-P, Millipore), using semi-dry method. Membranes were blocked for 2 hours at room tempertaure with 10% (w/v) dried non-fat milk powder in Tris-buffered saline with 0.1% Tween 20 (TBS-T). After blocking, the membranes were incubated at 4°C overnight with rabbit anti-BDNF antibody (Santa Cruz) diluted 1:500 in 5% (w/v) non-fat milk in TBS-T. Membranes were then incubated for 2 hours at room temperature with horseradish-peroxidase-labelled secondary antibody (Goat Anti-Rabbit IgG Antibody; Vector Laboratories) diluted 1:10,000 in 5% dried non-fat milk powder in TBS-T. After washing, the peroxidase activity was visualized on photographic film with ECL Prime Western Blotting Detection Reagent (Cytiva). The developed film was later digitized on ChemiDoc MP Imaging System (Bio-Rad) and analyzed using Image Lab software (Bio-Rad).

### Hippocampal organotypic culture and transfection

Hippocampal organotypic cultures were prepared from postnatal day 4-8 old mouse pups (either C57BL/6J or MMP-9 KO and their WT littermates). Hippocampi were dissected and sliced into 350 um thick transverse slices with a use of McIlwain tissue chopper and placed on PTFE membranes (Millicell; Merck, Cat# PICMORG50). Cultures ware kept in the air-medium interphase; medium composed of Minimal Essential Medium supplemented with, 20% heat inactivated horse serum, 12.9 mM D-Glucose, 5.2 mM NaHCO_3_, 30mM HEPES buffer, 2 mM MgSO_4_, 1 mM L-glutamate, 1 mM CaCl_2_, 0.075% Ascorbic Acid and 1 µg/ml insulin. Half of media volume was exchanged every 2-3 days. After 8-13 day in culture, slices were transfected using GeneGun (BioRad) method (Woods & Zito, 2008). Gene gun bullets were prepared with plasmids pCAG_GFP 30 μg DNA (structural change experiments); pCAG_MMP-9a1205cSEP and pCAG_mCherry ratio 1:1 50 μg DNA, or pCAG_MMP9a1205cGamillus with pCAG_mCherry ration 1:1, 50 μg DNA (MMP-9 release experiments). For experiments with blocked autocrine vesicular release – by co-expression of the tetanus toxin light chain (TeTx), pCAG_MMP9a1205cSEP was mixed with pCAG_mCherry-IRES-TeTx 1:1 50 μg of DNA; pCAG TrkB GFP and pCAG PLC RFP in ratio 1:3 50 ug DNA (TrkB activation experiment). Neurons expressing GFP and MMP-9 SEP/Gamillus with mCherry were imaged 1-7 days after transfection. Neurons expressing TrkB were imaged within 12-48h from the transfection.

### Two-photon FLIM

FRET measurements using a custom-built two-photon fluorescence lifetime imaging microscope was performed as previously described (Harward *et al*, 2016). GFP fluorophore was excited with 1.2-1.5 Mw, 920nm pulsating tunable laser (Ti-sapphire laser, Coherent, Cameleon). Emitted light was collected through 60x water immersion objective (NA 0.9, Olympus), split with a dichroic mirror (565 nm) and photons were detected by two separate photoelectron multiplier tubes (PMTs; H7422-40p, Hamamatsu). For green channel, light arriving to the PMT was filtered with a band-pass filter (510/70, Chroma) and for red channel using band-pass filter(620/90, Chroma). Fluorescence lifetime images were obtained using a time correlated single photon counting board (Time-harp 260, Pico-Quant) and processed by custom software FLIMage (https://github.com/ryoheiyasuda/FLIMage_public). Images for volume change measurements and TrkB activation were collected at 128×128 pixels with 7.8 Hz rate with 24 frame averaging. For MMP-9 SEP release imaging 64×64 pixels frames were collected with an 7.8 Hz rate with no averaging.

### Two-photon glutamate uncaging

A second Ti-sapphire tunable laser (Coherent, Cameleon) was set to 720 nm and used to uncage 4-Methoxy-7-nitroindolinyl-caged-L-glutamate (MNI-caged glutamate, Tocris). Up to four spines in separate regions of interest (ROI) per cell were stimulated simultaneously on four distinct secondary apical dendrites. Stimulation protocol consists of train of 6 ms, 2.7-3 mW pulses (30 times at 0.5 Hz) near a spines of interest.

Organotypic hippocampal slices were imaged in Mg^2+^-free ACSF solution (127 mM NaCl, 2.5 mM KCl, 4 mM CaCl_2_, 25 mM NaHCO_3_, 1.25 mM NaH_2_PO_4_ and 25 mM glucose) containing 1 µM tetrodotoxin (TTX, Tocris) and 2 mM MNI-caged L-gulatmate buffered with carbogen (5% CO_2_ and 95% O_2_). Experiments were performed at room temperature (around 24°C).

### Spine volume change measurements

Spines of GFP-expressing pyramidal neurons in CA1 subfield of hippocampus were imaged sequentially up to 4 spines located on separate secondary apical dendrites. For treatment, Slices were perfused in ACSF (see above) with either DMSO (for control; final concentration 0.08%), GM6001 (Cat: 364205, Calbiochem) in DMSO final concentration 25 μM or MMP-9/MMP-13 Inhibitor I (Cat: 444252, Calbiochem) in DMSO final concentration 5 μM. Experiments were repeated on organotypic cultures from at least three different dissections.

### Two-photon FLIM data analyses

The analysis of TrkB sensor activation was performed as previously described(Harward *et al*, 2016). To measure the fraction of a donor bound to an acceptor, we fit a fluorescence lifetime curve summing all pixels over a whole image with a double exponential function convolved with the Gaussian pulse response function:

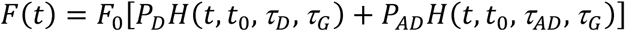

where τAD is the fluorescence lifetime of donor bound with acceptor, P_D_ and P_AD_ are the fraction of free donor and donor bound with acceptor, respectively, and H(t) is a fluorescence lifetime curve with a single exponential function convolved with the Gaussian pulse response function:

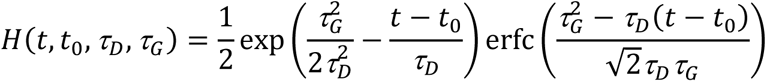

in which τ_D_ is the fluorescence lifetime of the free donor, τ_G_ is the width of the Guassian pulse response function, F_0_ is the peak fluorescence before convolution and t_0_ is the time offset, and erfc is the error function. We fixed τ_D_ to the fluorescence lifetime obtained from free eGFP (2.6 ns). To generate the fluorescence lifetime image, we calculated the mean photon arrival time, 〈t〉, in each pixel as:

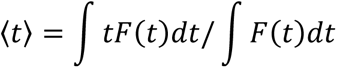

then, the mean photon arrival time is related to the mean fluorescence lifetime, 〈τ〉, by an offset arrival time, t_0_, which is obtained by fitting the whole image:

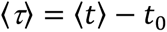

For small regions-of-interest (ROIs) in an image (spines or dendrites), we calculated the binding fraction (P_AD_) as:

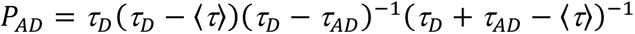

### TrkB activation

Images were collected simultaneously for up to 4 ROI located on distinct secondary apical dendrites of CA1 pyramidal neurons co-expressing TrkB-GFP and PLC-RFP constructs. Cells included in experiment had average initial binding fraction not higher than 45%. Slices were treated with either DMSO or Inhibitor I as described above. Experiments were repeated on organotypic cultures from at least three different dissections.

### MMP-9 release 2P imaging

Spines were imaged sequentially from distinct proximal secondary dendrites of pyramidal cells in CA1 co-expressing MMP9-SEP and mCherry or MMP-9 Gamillus and mCherry.

### Data analysis

Spine volume change was calculated using formula ΔF/F_0_ = (F-F_0_)/F_0_ where F is a sum of fluorescence in a given time from the ROI containing the spine of the interest and F_0_ represents average of F from the baseline of first 5-7 time points (before sLTP protocol).

MMP-9 SEP release was calculated using traces from 13 spines, where normalized intensity was calculated using formula ΔF/F_0_ = (F-F_0_)/F_0_ , where F_0_ corresponds to average of frame preceding the uncaging pulse and frame of uncaging pulse. Later the results from uncaging pulses 2-18 were averaged. Same analysis was performed for data collected from red channel corresponding to cytoplasmic mCherry treated as spine volume.

Binding fraction (P_AD_) was normalized by subtracting averaged baseline binding fraction (P_AD0_; before sLTP protocol) with a formula P_AD_ – P_AD0_.

## Acknowledgements

This study was supported by National Science Centre MAESTRO grant 2017/26/A/NZ3/00379 to L. K.;. D. L. was supported by Polish National Agency for Academic Exchange, Iwanowska Fellowship, NIH (R01MH080047, R35NS116804) to R.Y.

## Author contributions

Conceptualization, L.K. and P.M.; Methodology, R.Y.; Software, R.Y.; Investigation, D.L., B.K.; Writing – Original Draft, P.M.; Writing – Review & Editing, P.M., L.K., R.Y. and D.L; Visualization, P.M. and D.L.; Funding Acquisition, L.K., R.Y. and D.L.; Resources, L.K., K.K. and R.Y.; Supervision, P.M., L.K., K.K. and R.Y.

## Data Availability Section

This study includes no data deposited in external repositories.

---

**Supplementary Video 1. Spine-head enlargement during sLTP.**

Two-photon video demonstrating the volume increase upon uncaging during sLTP experiment. The images (maximal projections of Z-stacks) were acquired every minute before and after uncaging protocol. During the uncaging protocol (marked with appearing white dot) images were collected in a single plain with a 8 Hz frequency. Time is indicated on the video.

**Supplementary Video 2. MMP-9 release during sLTP.**

Two-photon video demonstrating the increase of MMP9-SEP fluorescence during the uncaging protocol. The video is in real time, images were acquired in a single plane at 7.8 Hz frequency. Uncaging pulses were delivered every 16 frames for 30 times. Time of a uncaging pulse can be visible as a red bar on the top for the frame. Red signal represents fluorescence of mCherry cytoplasmic fill and green signal represents MMP9-SEP.

**Supplementary Video 3. TrkB activation during sLTP.**

Two-photon FLIM video demonstrating activation of TrkB sensor. The images (maximal projections of Z-stacks) were acquired every minute before and after uncaging protocol. During the uncaging protocol (marked with appearing white dot) images were collected in a single plain with a 8 Hz frequency. Time is indicated on the video. Warmer colors represent shorter GFP lifetimes corresponding to increased binding fraction and higher TrkB activity.

